# Optimized chemogenomic library design strategies for precision oncology

**DOI:** 10.1101/2020.11.30.403329

**Authors:** Paschalis Athanasiadis, Balaguru Ravikumar, Neil O. Carragher, Paul A. Clemons, Timothy Johanssen, Daniel Ebner, Tero Aittokallio

## Abstract

Designing a focused compound screening library of bioactive small molecules is a challenging task since many compounds modulate their effects through multiple protein targets with various degrees of potency and selectivity. We describe here several analytic procedures with adjustable cut-off parameters that enable one to design anticancer target-focused compound libraries optimized for library size, cellular activity, biological and chemical diversity and target selectivity. Even though our focus was on designing compound libraries to enable a comprehensive investigation of the target biology of glioblastoma (GBM), the compound collections cover a wide range of protein targets and biological pathways implicated in various types of cancers, making the libraries widely applicable in precision oncology studies. We published the final screening set library, called the **C**omprehensive anti-**C**ancer small-**C**ompound **L**ibrary, or **C^3^L.**We hope these general library design principles and the current, widely annotated small molecule libraries will prove useful for the community in various phenotypic screening experiments in GBM and other cancers.

## Introduction

In the past ten years, there have been encouraging advances in the treatment and consequently, the survival rates, for many cancers. This progress has largely been driven by increased molecular understanding and classification of distinct cancer subtypes, the development of novel therapeutics focussing on targets associated with specific disease subtypes, and advances in both the type and range of therapeutic molecules available to oncologists for the treatment of patients. When considering molecules that have been approved or advanced in clinical trials, there are successful examples of targeted chemotherapeutics [1], small interfering RNAs [2], monoclonal antibodies [3], micro-RNAs [4] and virotherapy [5], to name a few. These developments are very encouraging considering the complexity of human cancers and the difficulty in developing effective treatments for the advanced disease. However, small compound chemotherapeutics still makeup the vast majority of approved drugs available to oncologists treating cancers [6–8]. For example, in the case of glioblastoma (GBM) brain tumours, small molecule chemotherapeutics are currently the only approved treatment modalities beyond surgery and radiation [9], and represent the most fertile ground for future innovation of new therapeutics which address the severe limitations inherent in treating brain-tumours. These limitations include (i) breaching the blood-brain barrier to effectively deliver therapeutics to the tumour, (ii) developing combinatorial treatments to systematically target multiple pathway redundancies and tumour vulnerabilities inherent in brain tumours which typically exhibit dynamic intra and inter tumour heterogeneity, and (iii) selectively targeting glioblastoma stem cells, which have been shown to be the main source for cancer recurrence in GBM [10].

Traditional drug development often employs high-throughput drug screening of very large collections of diverse small molecule libraries against a nominated therapeutic target to identify chemical starting points for further development. This process has been relatively successful at the industrial level, but it is generally less successful at the academic level due to the prohibitive infrastructural costs required to develop and produce larger screens, as well as the cost to develop a small compound hit into a clinical candidate drug. Instead, many academic target discovery and drug development facilities have focussed on developing more physiologically relevant phenotypic assays, which better recapitulate key areas of disease biology [11], and screening of smaller, more focussed libraries of small compounds, such as collections developed and curated for drug repurposing [12–14], probes for target discovery [6, 15], or pharmacologically active small compound sets [7]. This focussed approach has the advantage of providing a better understanding of the molecular basis of the disease, while simultaneously providing the opportunity to exploit existing therapeutics and compounds with known safety profiles, along with probes that possess drug-like properties and compounds with known protein targets. In complex diseases of unmet therapeutic need where target biology is poorly understood or where disease heterogeneity indicates multiple target pathways contribute to disease progression (as exemplified by GBM) phenotypic screening of target-annotated compound libraries in relevant patient-derived cell models provide a valuable strategy for empirical identification of drug targets or drug combinations which address such disease complexity. By circumventing major pitfalls and hurdles, such as poor selectivity, cellular activity and biological/target space diversity, these libraries have the potential to greatly accelerate the drug development process.

Here, we describe the analytical steps for the construction of a new, comprehensive anti-cancer target-annotated compound library, designed to interrogate a wide range of potential cancer targets in phenotypic screening. The library design is approached as a multi-objective optimization problem, where the aim is to maximize the cancer target coverage, while maintaining compound cellular potency and selectivity, and minimizing the number and cost of the compounds arrayed into the final screening library. To do this, we used two target-based approaches. First, we defined all of the proteins implicated in cancers and searched for small-molecules against the druggable cancer targets among approved and investigational compounds identified from the literature, existing oncology collections, and compounds identified through manually searching of clinical trials databases. Second, to expand the target-annotated compound library, we surveyed several pan-cancer studies to systematically identify anti-cancer compound-target pairs, and then expanded the chemical space around those novel targets by identifying additional bioactive compound probes through database queries. Importantly, cancer-mutated proteins, first neighbors and influencer targets were further investigated for potential small compound interactors, which then collectively generated a large *in-silico* probe set collection. Finally, we refined the probe set collection by applying several filters, including optimized activity and similarity thresholds, and removal of redundant structures and compounds that could not be readily sourced, to yield a sufficiently diverse, focused, and annotated compound library for phenotypic screening purposes. We have published both the approved and investigational compound collection as well as the final probe collection as datasheets, which together form the **C**omprehensive anti-**C**ancer small-**C**ompound **L**ibrary, or **C**^3^L.

## Results

### Defining and collating a comprehensive list of cancer associated protein targets

Our first design objective was to define a comprehensive list of protein targets associated with the development and progression of cancers, which could then be employed to form the basis of the anti-cancer small-molecule library. Our target space was designed to span a wide range of protein families, cellular functions and cancer types, and covers all of the divisions of the ‘hallmarks of cancer’ [16]. We first defined a list of proteins known to be implicated in cancers using The Human Protein Atlas [17] and nominal targets of pan-cancer studies from the PharmacoDB [18], covering the target space of 946 oncoproteins (see Methods for details).

### Identifying and curating small-molecule inhibitors of cancer associated targets

After defining the comprehensive list of cancer-associated protein targets, our next objective was to identify and curate a small-molecule collection of compounds targeting these proteins. Since the sources of the compounds and targets ranged from investigational and experimental probe compounds to the approved drugs, we took a systematic approach to defining each source, sorting the compounds targeting the cancer targets, ranking the compounds for activity, diversity and availability for sale, and finally, producing a sortable and searchable database defining the screening library consisting of two separate compound collections described below.

### Compound set 1 - Experimental probe compound (EPC) collection

In the EPC collection, compound-target pairs were extracted manually from public databases. The experimental probe compound collection contains chemical probes and investigational compounds from three nested subsets of decreasing sizes; (i) the **theoretical set** which is an *in-silico* set curated from established target-compound pairs for a broad target space (1655 proteins), (ii) the **large-scale set** is a large-scale screening collection of filtered compounds covering the same target space, and (iii) the **screening set** is the final set of most potent probes arrayed into the physical library. The screening set is also the smallest subset due to the limitations in compound availability for sale for screening purposes. A schematic diagram of the construction of these three compound sets is illustrated in **Figure 1**, and described in more detail in the following subsections (see Methods for the step-by-step procedures).

**Figure 1.**
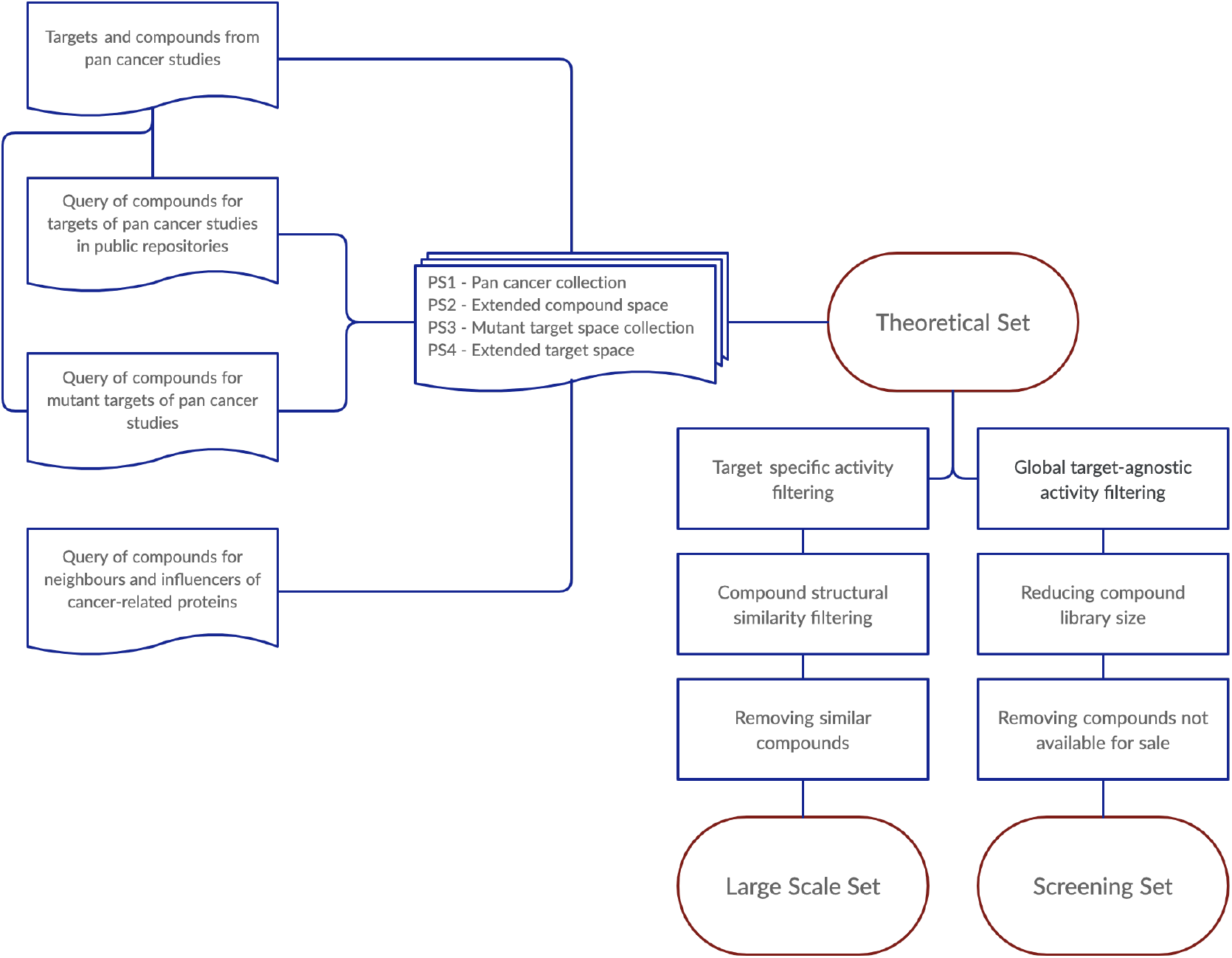
Workflow of acquiring the three probe compounds sets (Theoretical, Large-scale and Screening set). Four probe sets (PS) were defined: The pan-cancer collection (PS1) includes compounds and their annotated nominal targets from various pan-cancer studies. The extended compound space (PS2) consists of compounds that have off-target activity against the annotated targets of the pan-cancer studies, while excluding compounds already in the PS1 set, retrieved from public drug/target repositories such as ChEMBL [19], Drug Target Commons (DTC) [20] and DrugBank [17]. The collection for the mutant target space (PS3) consists of compounds that have activity against the mutant variants of the annotated targets extracted by using the COSMIC database [21]. The extended target space collection (PS4) extends the targets space of cancer-related targets through the nearest neighbor approach [22]. See Methods for the details of the construction of the various probe collections.

**Theoretical set** contains 336,758 unique compounds from four probe sets (PS): pan-cancer collection, pan-cancer collection with extended compound space, mutant target collection, and mutant target collection with extended compound space (see Methods for details). **Large-scale set** contains a subset of compounds from the Theoretical set, filtered to reduce the number of molecules in the library while covering the same target space (**Figure 2**), using both the activity and similarity filtering procedures with pre-defined cut-off values (note: the cut-off parameters are freely adjustable in other studies). The large-scale set was based on both the on- and off-target profiles of the small-molecule compounds (n=2282), which could be used in larger-scale screenings campaigns in academic or industrial projects (**Table 1**).

**Figure 2.**
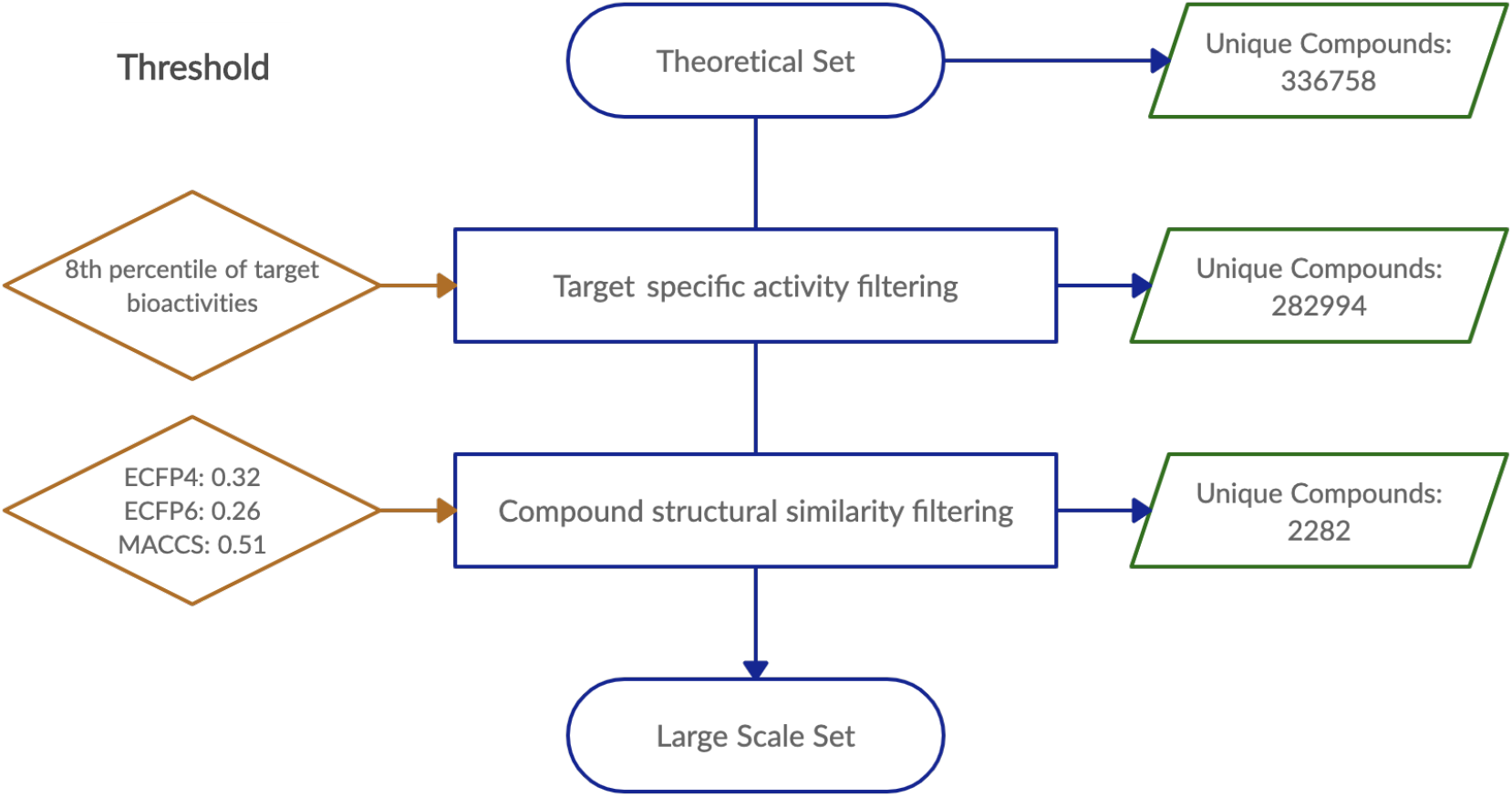
The procedure for the design of the Large-scale probe compound set. The target-specific activity filtering is based on either biochemical or cell-based evidence that the compound has a potency against the particular target (less stringent filtering). Compounds that show sufficient similarity to other compounds based on two fingerprints out of the three were removed in the structural similarity filtering step (more stringent filtering). The thresholds used in the current design procedures are shown on the left. See Methods for details of the filtering procedures.

**Table 1.** The compound and target spaces of the three probe compound collections.

| Probe collection | Unique compounds | Unique protein targets | Unique interactions* |
| --- | --- | --- | --- |
| Theoretical set | 336758 | 1655 | 494320 |
| Large-scale set | 2285 | 1655 | 8137 |
| Screening set | 1211 | 1386 | 2699 |

\* Unique compound-target activities by considering various bioactivity types from multi-dose assays as repeated measurements of the same compound-target interaction.

**Experimental screening set** contains a smaller number of 1211 compounds that can be used for screening applications, thus making the probe set suitable for routine exploration of oncology-associated biological target space in complex phenotypic assays and identification of potential candidates for further drug development. This probe subset was obtained by subjecting the Theoretical set to three filtering procedures to obtain a manageable library size for screening (see **Figure 1**). These procedures with freely adjustable parameters involve (i) global target-agnostic activity filtering to remove non-active probes, (ii) selecting the most potent compound for each target to reduce the compound space, and (iii) checking which compounds are available for sale (see Methods for details). Since two of the Theoretical sets are already small in size (PS1 and PS3), the global activity filtering was implemented on the two larger subsets only (PS2 and PS4).

### Compound set 2 - Approved and investigational compound (AIC) collection

According to its design principles, the above-described probe compound set included mostly compounds that are currently in pre-clinical stages (**Figure 3A**). We therefore next searched for additional small-molecules that are currently either approved for chemotherapeutic or immunotherapeutic treatments, or anti-cancer compounds in clinical development stage. This compound collection was manually curated from several public resources (**Table 2)**, and it partly overlapped with the probe set collection (**Figure 3C**).

**Figure 3.**
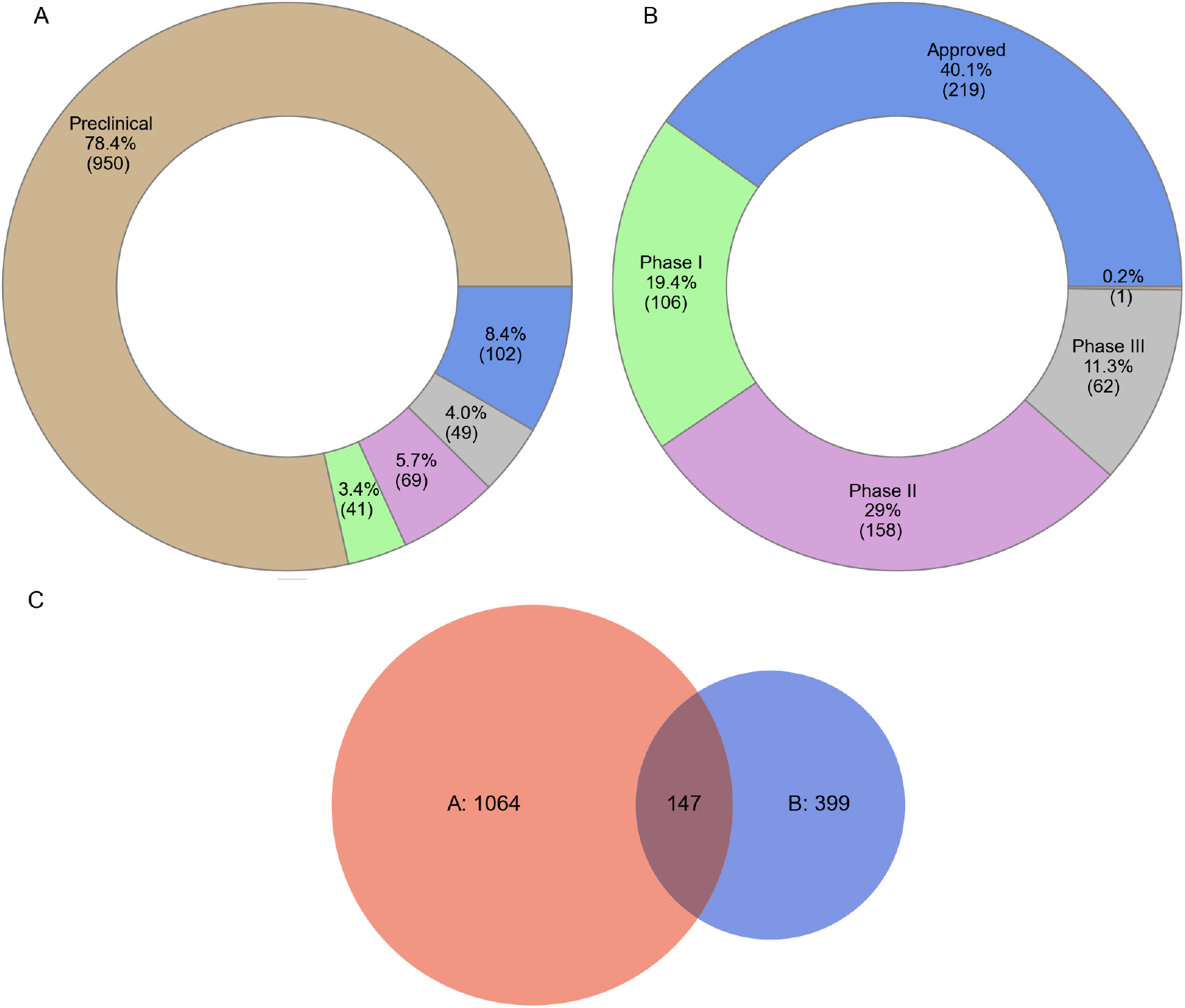
Clinical development phase distribution of the compounds in the two collections. (A) experimental probe compound (EPC) collection, and (B) approved and investigational compound (AIC) collection. The clinical development stage was extracted from ChEMBL [19]. Numbers in parentheses indicate the number of compounds in each category. (C) Overlap of the compounds between the EPC and AIC collections. The numbers in the Venn diagram indicate the numbers of unique and shared compounds in the two sets, respectively.

**Table 2.** Resources used to prepare the approved and investigational compound collection.

| Resource | Number of compounds |
| --- | --- |
| Approved kinase inhibitors<sup>a</sup> | 62 |
| FIMM oncology collection<sup>b</sup> | 460 |
| Glioblastoma therapeutics<sup>c</sup> | 25 |

<sup>a</sup>FDA-approved kinase inhibitors retrieved manually from a recent review article [23].
<sup>b</sup>Manual curation of small-molecules for their approval status in different countries; FIMM, Institute for Molecular Medicine Finland (FIMM).
<sup>c</sup>Approved and investigational compounds for GBM retrieved by manually searching Clinical Trials database (https://www.clinicaltrials.gov/) and SynergySeq platform [24].

The AIC collection was further subjected to removal of duplicates and similarity searches using the ECFP4, ECFP6 and MACCS fingerprints (see Methods), where a similarity threshold of ≥0.99 was used to identify and remove highly similar compounds (e.g. doxorubicin and epirubicin, with the first compound being removed from the final set). Following the above curation procedure, the final AIC collection consisted of 546 unique compounds. The distribution across different clinical development phases is shown in **Figure 3B**.

### Characterization of the compound and target spaces of the probe collections

We next analyzed the compound and target spaces of the probe compound collections, designed using the above-described procedures and the selected parameter values for filtering. The target distributions of the three probe collections remained relatively similar, where the screening set shows reduced numbers of multi-target compounds (**Figure 4A**). However, the median number of targets per compound was one for each probe collection, indicating that the sets include relatively selective compounds. For most of the targets, the screening set contains only a single compound per target, whereas in the Theoretical and Large-scale sets the median number of compounds per target was 42 and 2, respectively (**Figure 4B**).

**Figure 4.**
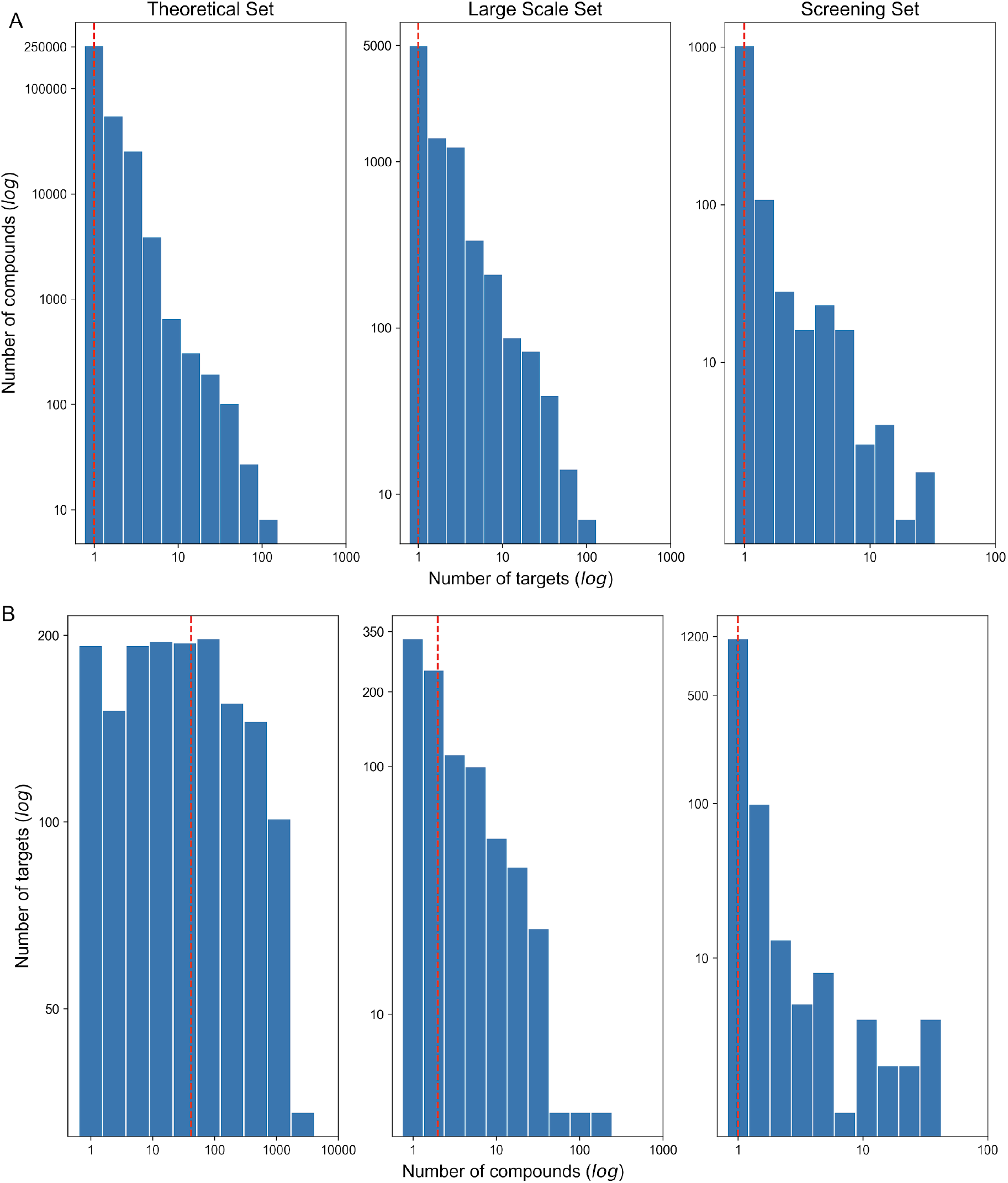
(A) The number of targets per compound, and (B) the number of compounds per target in the three compound collections. The dashed line indicates the median. The x-axis and y-axis are log10-scaled in each panel, while the numbers present the non-logged counts (note the different scales between the panels). The counts are based on the target activity threshold ≤1000nM (see **Suppl. Fig. 1** for other activity thresholds).

The target classes of the screening set represent well those of the larger probe collections covered in the Theoretical and Large-scale sets (**Figure 5, Suppl. Fig. 2**). As expected, inhibitors for kinases and other enzymes are the most frequent in the three probe collections, but there are also other well-covered target classes such as membrane receptors, epigenetic regulators and ion channels. Among the kinase inhibitors, the screening probe set covers all the major kinase families, and only a few of the kinase targets present in the Theoretical/Large-scale sets are missed in the smaller Screening set (**Figure 6**). Importantly for anticancer applications, the compound collections cover many targets implicated in various types of cancers, with breast and lung cancers having largest numbers of targeted compounds followed by glioma (**Figure 7**).

**Figure 5.**
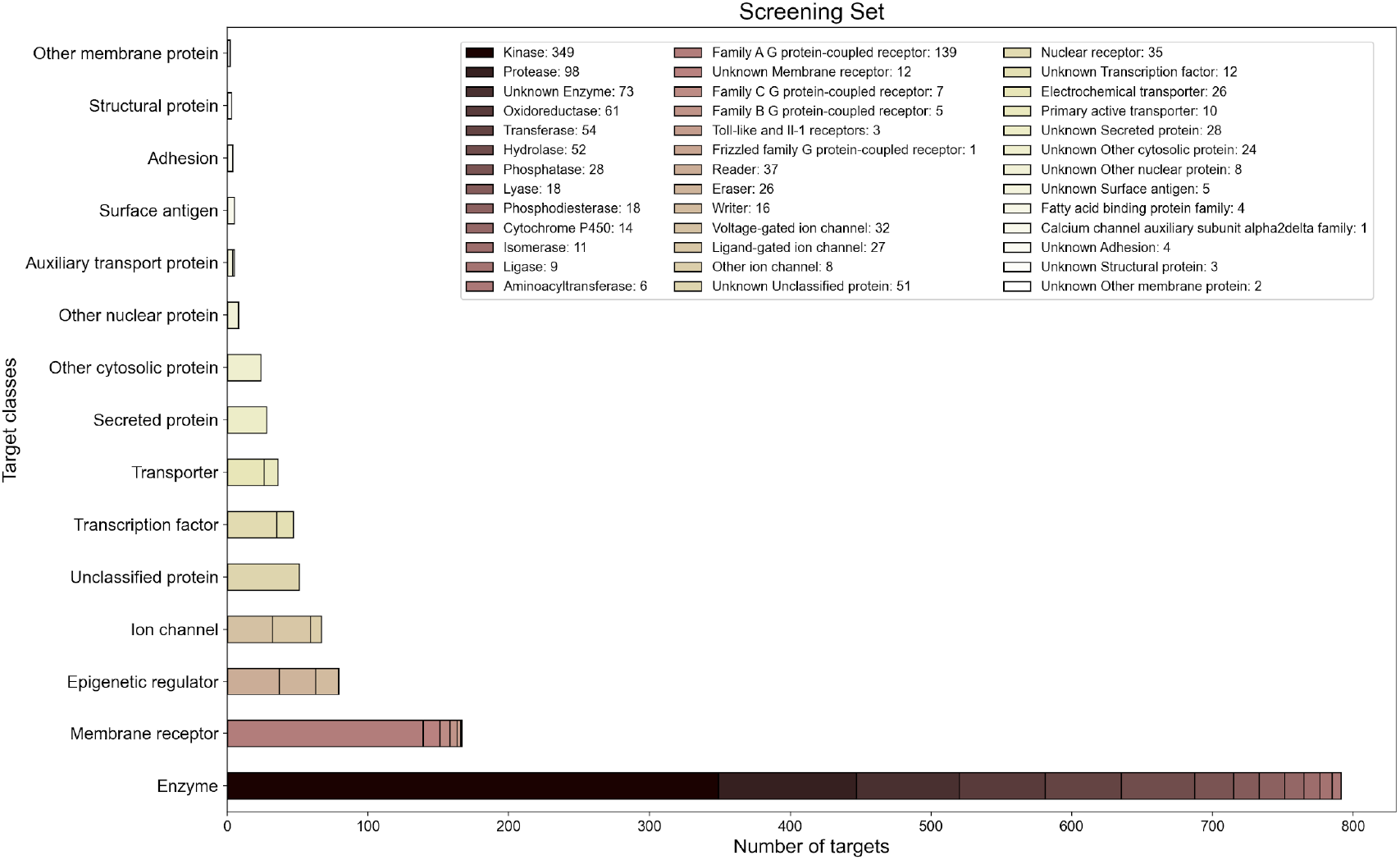
The distribution of target classes of compounds in the Screening set. For the Theoretical and Large-scale sets, **see Suppl. Fig. 2**. The target classification of proteins was extracted from ChEMBL [19].

**Figure 6.**
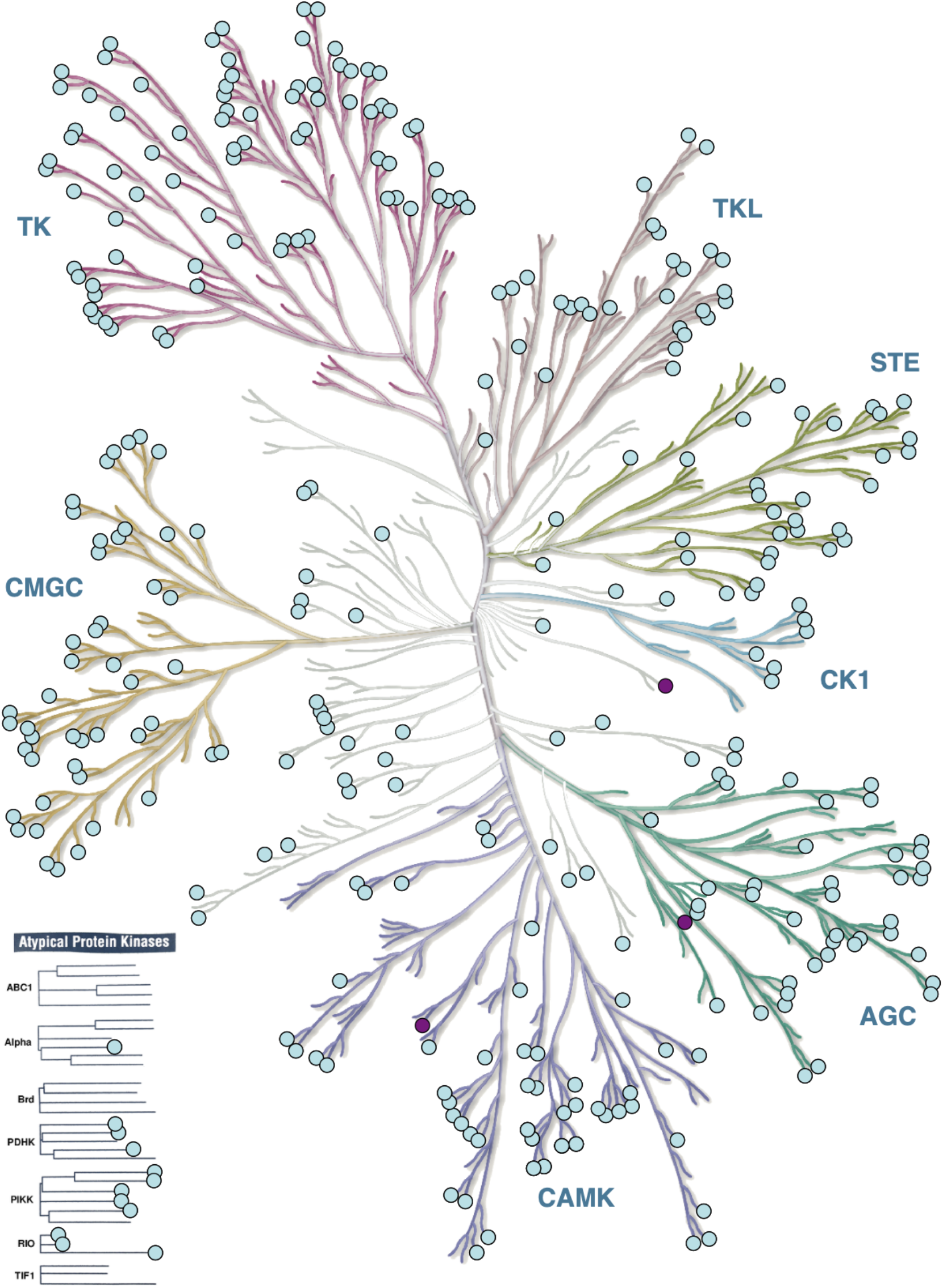
Kinase families covered by the targets of the Theoretical/Large Scale set and Screening set. The kinases colored in purple indicate the targets present only in the Theoretical/Large-scale sets, but not in the Screening set, while the light-green targets are covered by all the three collections. KinMap[25] web-tool was used for the creation of the illustration, reproduced courtesy of Cell Signalling Technology, Inc. (www.cellsignal.com).

**Figure 7.**
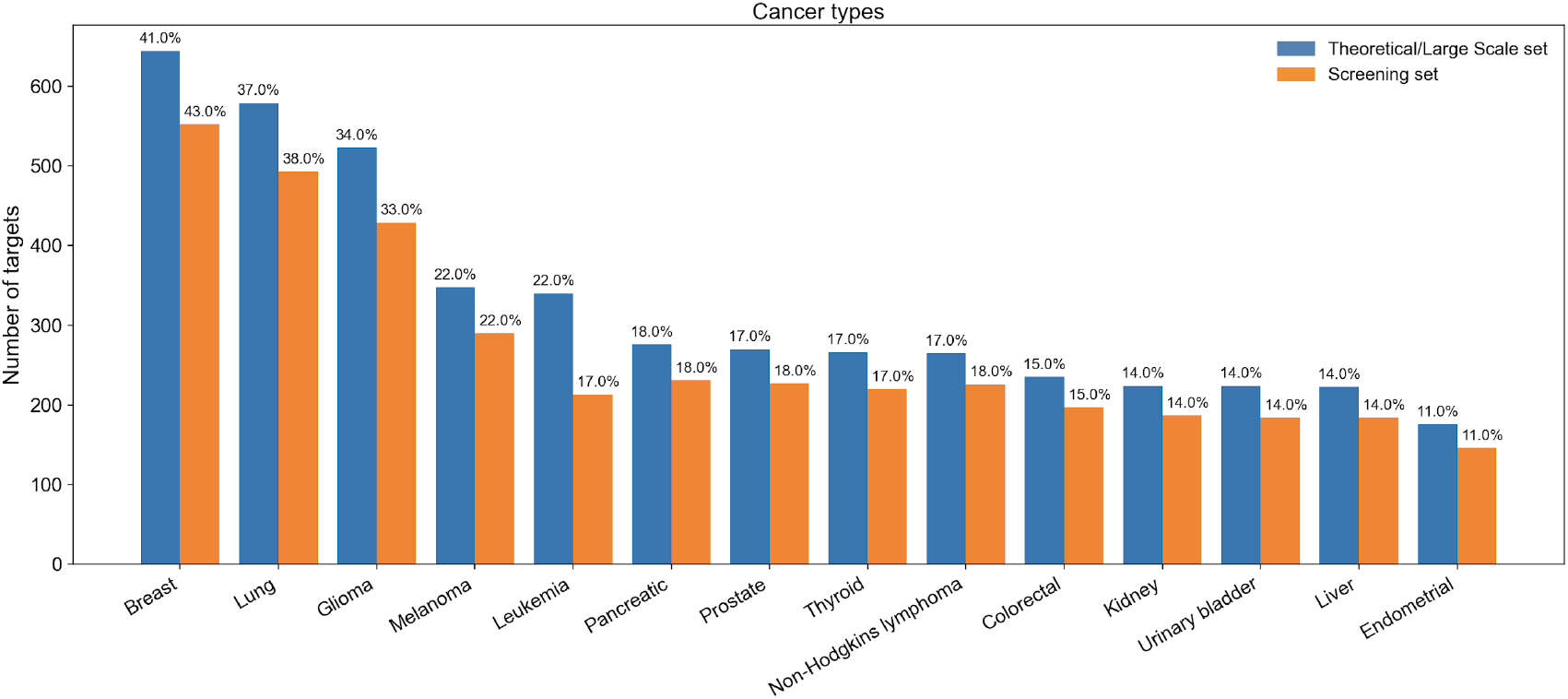
The number of targets associated with cancer types in Theoretical/Large-scale and Screening sets. The numbers above the bars indicate the percentage of targets in the two collections of different sizes. The disease associations were extracted from OpenTargets [26], with overall association score >0.5 (https://www.opentargets.org/).

## Methods

### 1. Approved and investigational compound (AIC) collection construction

The selection and the curation of compounds in the AIC Collection was carried out manually. The compounds’ IDs (from PubChem[27] and ChEMBL[19]) and their structural description (canonical SMILES) were retrieved using PostgreSQL [28]. One compound (TAK-530) was found to have neither a compound ID nor a canonical SMILES, and it was therefore removed. The search for approved compounds for GBM was carried out by manually searching the Clinical Trials database (https://www.clinicaltrials.gov/), wherein compounds currently approved or clinically investigated for GBM were found, and the availability of these compounds were later determined by crosschecking the compound in the SelleckChem (https://www.selleckchem.com/) and PubChem (https://pubchem.ncbi.nlm.nih.gov/). The fingerprint descriptors of compounds were enumerated using the RDKit [2] chemoinformatics module in Python 3.7 [30]. A similarity threshold of ≥0.99 was used to identify and remove highly 2 similar compounds (doxorubicin and epirubicin, with the first compound being removed). The resulting AIC collection consisting of 546 unique compounds is available in GitHub (https://github.com/PaschalisAthan/C3L).

### 2. Experimental probe compound (EPC) collection construction

In the EPC collection, compound-target pairs were extracted manually from pan-cancer studies using public databases (PharmacoDB [18] and The Human Protein Atlas [31]). The wild type and mutant variants of the targets, along with the first neighbors and influencers of other cancer-related targets [22], were later used to search for additional compounds that demonstrated sufficient target activity using PostgreSQL [28] (see below subsections).

Next, analytical compound filtering procedures were used to produce the three probe collections (the Theoretical, the Large Scale and the Screening set) using RDKit [29], where the procedures involved checking the structural similarity between the compounds, and Python scripts for additional processing, such as adding extra annotations for the targets or the compounds extracted. The full list of cancer associated protein targets and associated probe compounds can be found at https://github.com/PaschalisAthan/C3L.

#### 2.1. Pan-cancer probe collection - PS1

This collection of probe compounds focuses on compounds and targets implicated in various types of cancers. To construct this collection, several comprehensive large-scale pan-cancer studies were analyzed (see **Table 2.1**). A set of 946 unique targets and 1525 compounds were curated from pan-cancer studies using the nominal targets from the PharmacoDB database [18], and The Human Protein Atlas database [31]. After removing redundant compounds, a total of 851 unique compounds remained that cover the target space of 946 unique proteins.

**Table 2.1.**
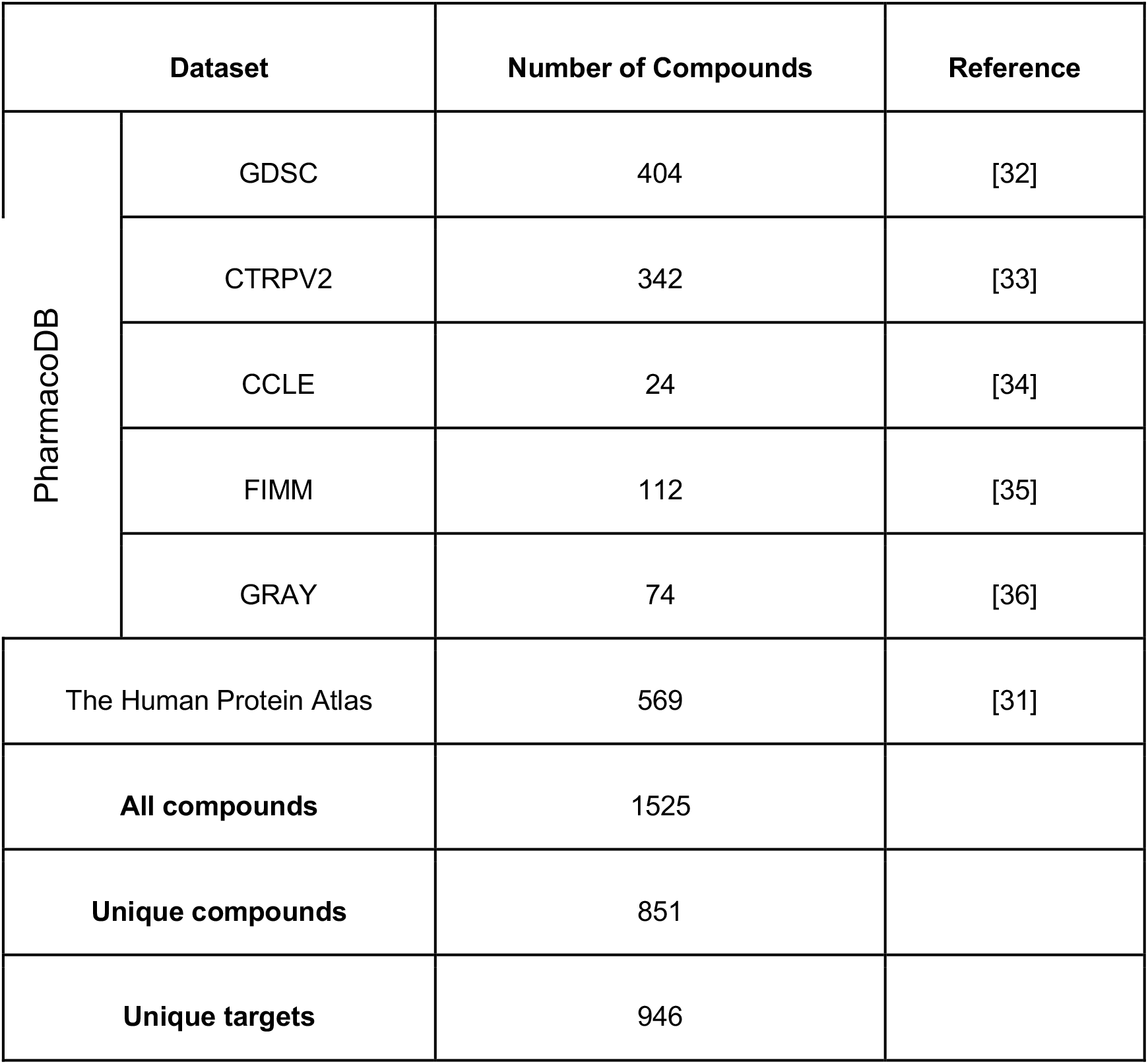
The datasets listing the intended targets implicated in pan-cancer studies.

Since the degree of the cellular activity of the compound-target interactions was needed in the activity filtering and extension of the target profiles to include also potent off-targets, the preferred name of the compounds was mapped to ChEMBL IDs, along with the targets Uniprot IDs to find the reported compound-target interaction potencies either in ChEMBL [19], DrugTargetCommons [20] or DrugBank [17] databases. During this process, 285 additional interactions between 185 compounds and 130 targets were found in these databases using multi-dose assays (IC_50_, EC_5_, K_d_ and K_i_).

#### 2.2. Extending the compound space - PS2

To further extend the compound space, the annotated primary targets from the pan-cancer studies were queried across publicly available compound bioactivity repositories [17, 19, 20]. Using a relatively liberal activity threshold of <= 1000 nM and multi-dose activity types, such as IC_50_, EC_50_, K_i_ or K_d_, the compounds having cellular activity against these targets were curated. In case of multiple entries corresponding to the same compound-target interaction, the median activity value was recorded [37]. This curation procedure resulted in 141,087 unique compounds (**Table 2.2**).

**Table 2.2.** Number of compounds that show activity agents the pan-cancer targets.

| Repository | Number of Compounds | Reference |
| --- | --- | --- |
| ChEMBL | 140 626 | [19] |
| DTC | 99 565 | [20] |
| DrugBank | 845 | [17] |
| **All compounds** | 241 036 |  |
| **Unique compounds** | 141 087 |  |

#### 2.3. Collection for the mutant target space - PS3

In addition to investigating the wild type targets implicated in various cancers (Section 2.2), their corresponding mutant variants were also analyzed. The mutation information of the annotated targets of Section 2.1 were retrieved from the COSMIC Database [21]. The various types of variants included in the current curation process are listed below:

- Substitution - missense
- Substitution - coding silent
- Complex - compound substitution
- Insertion - in frame
- Complex - deletion in frame
- Substitution - nonsense
- Deletion - in frame

These mutant targets were queried across the existing data repositories ChEMBL [19] and DTC [20], and the corresponding compounds were compiled using the criteria similar to those described in Section 2.2. The resulting collection consists of 944 unique compounds targeting 293 unique mutant targets (**Table 2.3)**.

**Table 2.3.** The number of compounds that show activity against the mutant variants.

| Repository | Number of Compounds | Reference |
| --- | --- | --- |
| ChEMBL | 944 | [19] |
| DTC | 446 | [20] |
| Unique compounds | 944 |  |

#### 2.4. Extending the target space - PS4

Finally, as an effort to extend the target space, we queried public repositories for cancer-related targets, their first neighbors and influencers in four cancer types with high mortality rate (colon, breast, hepatocellular, non-small cell lung cancer). Such extended “targets” are suggested to influence cancer pathogenesis and therefore to increase the drug target space for anticancer therapies [22]. A target was considered as a cancer-related gene, when the corresponding protein was either mutated or had a differential expression in cancer. A first neighbour of cancer-related protein was defined as a protein that is directly and physically interacting with a cancer-related protein in human interactome or signalling networks [22], according to the databases SignaLink 2 [38], Reactome [39], HPRD [40], DIP [41], IntAct [42], BioGrid [43], or in a cancer signalling network [44]. An influencer protein was defined as a protein that has a direct interaction to one of the first neighbours. After combining these three subsets, an overlap analysis with the cancer-related targets was performed. The non-overlapping targets were queried in ChEMBL [19] and DTC [20] to find a total of 208 653 potent compounds following the same procedure as in the previous sections (**Table 2.4)**.

**Table 2.4.**
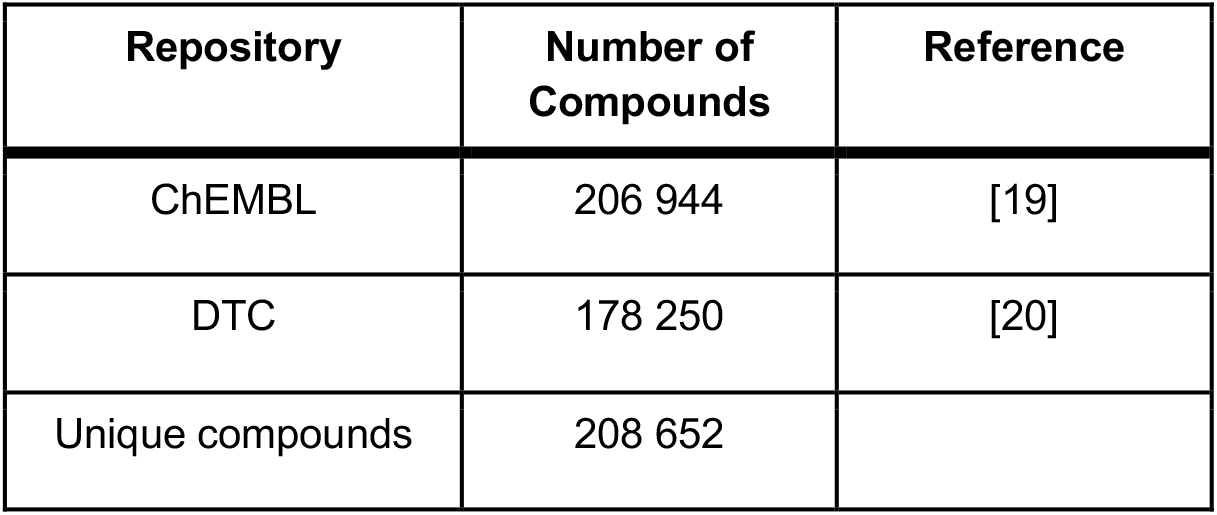
Number of compounds that show activity against the extended target space.

| Repository | Number of Compounds | Reference |
| --- | --- | --- |
| ChEMBL | 206 944 | [19] |
| DTC | 178 250 | [20] |
| Unique compounds | 208 652 |  |

### 3. Reducing the number of compounds

To make the Theoretical screening set that consists currently of 336758 probes more feasible for academic screening projects, several filtering procedures were subsequently applied, each with freely adjustable cut-off parameters that determine the stringency of the filtering process, and hence the number of compounds that pass it.

### 3.1. Target-specific activity filtering

The first library filtering technique was to apply a target-specific activity threshold to reduce the number of compounds from the Theoretical set. More specifically, the procedure comprised of the following steps:

1. All the activity values were log-transformed
2. Repeat for each target:

1. Target’s activity distribution was normalized to zero mean and unit variance
2. A threshold was selected such that 80% of the target’s activities (i.e 80% percentile) remain within that threshold (see an example in **Table 3.1**)
3. Compounds with activities higher than the selected threshold for the target were removed (i.e. these show less potency toward the particular target)

**Table 3.1.** An example of how the activity threshold was selected for the target CIT (Citron Rho-interacting kinase - O14578) based on Kd multi-dose bioactivity readouts.

| Target | Compound | Activity | Value (nM) * |
| --- | --- | --- | --- |
| O14578 | CHEMBL379218 | Kd | 13 |
| O14578 | CHEMBL574738 | Kd | 52 |
| O14578 | CHEMBL603469 | Kd | 85 |
| O14578 | CHEMBL558752 | Kd | 87 |
| O14578 | CHEMBL1721885 | Kd | 94 |
| O14578 | CHEMBL428690 | Kd | 110 |
| O14578 | CHEMBL1287853 | Kd | 140 |
| O14578 | CHEMBL494089 | Kd | 200 |
| O14578 | CHEMBL572881 | Kd | 300 |
| O14578 | CHEMBL607707 | Kd | 310 |
| O14578 | CHEMBL388978 | Kd | 340 |
| O14578 | CHEMBL10 | Kd | 420 |
| O14578 | CHEMBL475251 | Kd | 440 (Percentile 80%) |
| O14578 | CHEMBL278041 | Kd | 510 |
| O14578 | CHEMBL1908397 | Kd | 650 |
| O14578 | CHEMBL553 | Kd | 680 |

\*Cells in green are the activities that are retained after applying the target-specific activity filtering. The bioactivities are arranged from most potent (small Kd) to less-potent (high Kd).

#### 3.2. Compound structural similarity filtering

The next filtering step was to find the similarity threshold above which two compounds were considered sufficiently similar. This procedure was based on the assumption that similar compounds are expected to have similar activity distributions. The cut-off value for the similarity was identified with the Akaike Information Criterion, which is an estimator for the degree of information that is lost when using a candidate model, i.e., smaller values indicate low information loss that is better than high values indicating high information loss [45]. More specifically, the procedure for defining the similarity cut-off used the following steps:

1. A portion (here, 10%) of the total number of compounds was chosen for the similarity cut-off estimation
2. Each compound’s similarity (based on ECFP4, ECFP6 and MACCS fingerprints) was estimated with the rest of the compounds within the portion, hence leading to a compound similarity matrix
3. A similarity threshold was varied, from 0.1 to 0.99 (with 0.01 step size), and for each threshold:

1. Compounds that had similarity value equal or greater than the threshold were identified
2. The similarity distributions of those compounds were compared using the Kolmogorov-Smirnov (K-S) test and the test statistics were recorded
3. The Akaike Information Criterion (AIC) value was calculated based on the K-S test statistic values
4. The optimal similarity threshold was selected based on the smallest AIC value (see **Figure 3. 2**)
5. Steps 1-4 were repeated thrice using 3 different random seeds

**Figure 3.2.**
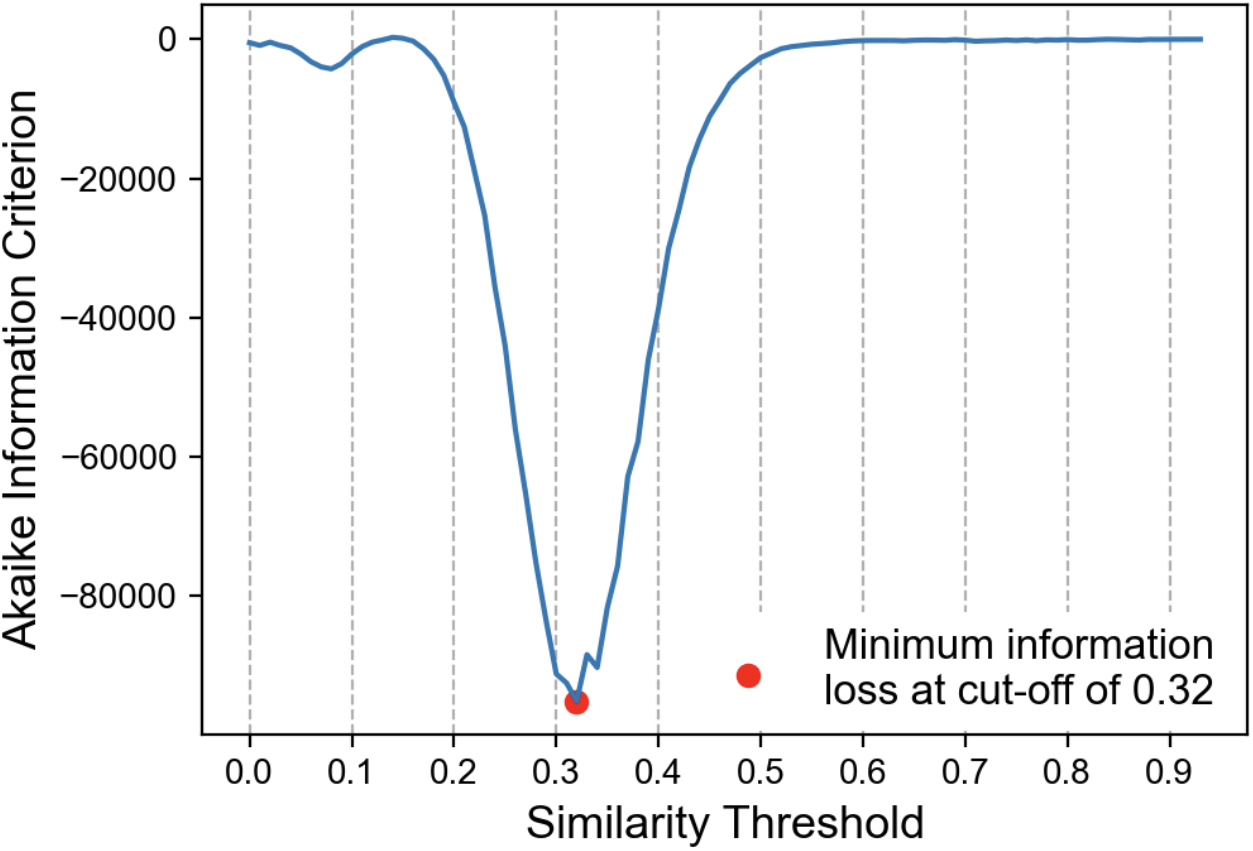
An example how the optimal similarity threshold was identified based on the Akaike Information Criterion. The optimal structural similarity threshold was defined for each structural fingerprint separately as the smallest value of the curve (thresholds ranging from 0 to 1), indicating the lowest information loss.

#### 3.3. Global activity filtering

The first step for reducing the size of the physical screening library was to apply an activity threshold similar to that explained in Section 3.1, but instead of being target-specific as above, herein the same activity threshold was used across all the targets (i.e., target-agnostic, global filtering). More specifically, the procedure of applying a global activity filtering followed these steps:

1. The profile of quantitative bioactivity values for each target were recorded when creating the theoretical set (see Section 2.1)
2. The bioactivity values are log-transformed and normalized to zero mean
3. The target with the highest activity standard deviation is identified
4. An activity threshold is selected such that 95% of the bioactivities of the targets from step 3 are within the selected threshold (see **Figure 3.3**).
5. The selected activity threshold is applied to all the targets to remove the compounds with bioactivities larger (i.e. less potent) that the selected threshold

**Figure 3.3.**
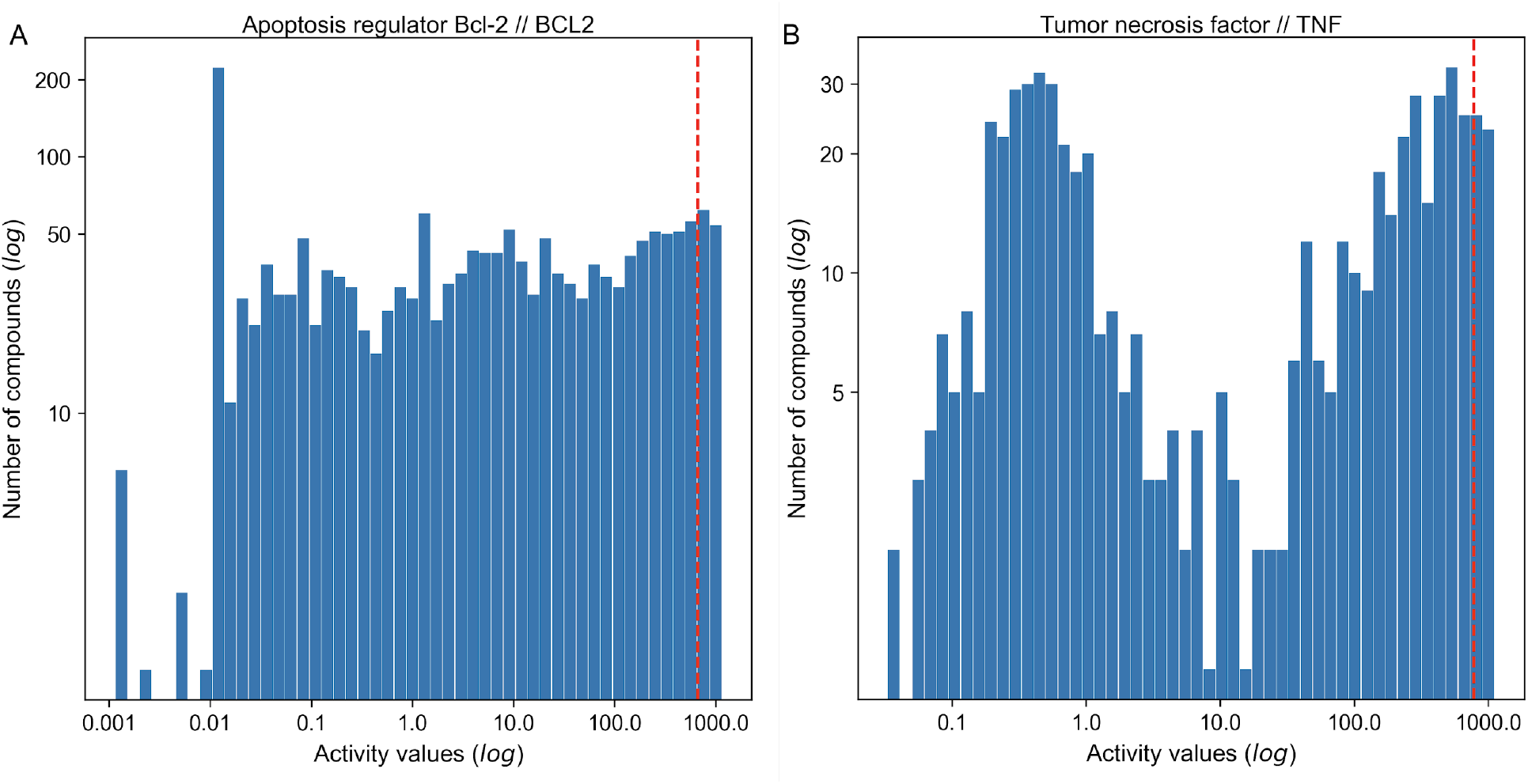
Examples of protein targets with the highest standard deviation of compound bioactivities based on which the global activity threshold was selected for the 4 probe sets. (A) Activity threshold of 670nM (red dotted line) in the PS2, and (B) activity threshold of 780nM (red dotted line) in the PS4.

#### 3.4. Reducing the compound space

The second step for the screening library size reduction was to pick only a single compound for each target to have the smallest number of most potent compounds for covering the same target space. More specifically, the compound that had the lowest activity value among the multi-dose activity types (IC_50_, EC_50_, K_i_, K_d_) was selected for the particular target, since that compound is assumed to have the highest binding potency against the target. The different activity types were treated equally in this process.

#### 3.5. Compound availability filtering

The final step for designing the screening library was to include only the compounds that are for sale from at least one vendor using availability information for ZINC15 [46]. More specifically, if a particular compound was not available for sale, then the next most potent compound that is available for sale replaced the original one in the screening library. Even though this step reduced the number of compounds in the final screening library to 52% of the original size, the targets’ coverage remained at 86% of the original space (**Table 3.5**). Furthermore, the potency distributions remained relatively unchanged (**Suppl. Figure 3**), where the differences originated mainly from the few of the most potent compounds.

**Table 3.5.** The compound and targets spaces of the screening set before and after replacing the unavailable compounds.

| Probe set | Unique Compounds | Compounds for-sale* | Unique targets | Unique interactions |
| --- | --- | --- | --- | --- |
| Screening set initial | 2331 | 635 | 1628 | 5315 |
| Screening set after compounds replaced | 1211 | 1211 | 1386 | 2699 |

\* The availability for sale information is extracted from ZINC15 [46].

## Discussion

In the field of precision oncology, there have been several efforts to systematically catalogue all of the genes implicated in cancer [47, 48]. Several commercial and governmental chemical genomics and approved drug libraries targeting many of the most heavily investigated genes are readily available [12, 13]. However, even though these commercial or in-house libraries have been well-utilized in many anticancer screens and drug repurposing studies [14,15, 49,50], there is a need for a comprehensive, target-annotated, anti-cancer library optimized for selectivity and diversity against all known oncology targets. There are well-annotated libraries for specific classes or applications, such as for kinase inhibitors [51] and drug repurposing [52], respectively, but what is lacking is a comprehensive probe library that provides a starting point for various anticancer screening applications. Despite the emergence of new treatment modalities, such as monoclonal antibodies, molecularly-targeted small compounds are, and will most likely remain, the most prolific anticancer therapeutics for the foreseeable future that are also applicable in advanced stages of the disease, for instance, after tumour has progressed towards metastatic disease and when the cancer has become chemo-or radio-therapy resistant [6–8]. The development and implementation of a well curated and comprehensive screening library will facilitate the identification of novel anticancer drug targets, drug combinations, synthetic-lethal interactions and novel targets to overcome cancer resistance. Subsequent application may include the identification of chemical starting points, new target-directed drug discovery programs and approval of new therapeutics in specific indications

We formulated the compound library construction as a multi-objective optimization (MOP) problem, where the aim is to simultaneously maximize the anticancer target diversity and compound potency (or selectivity), while minimizing the number and cost of the compounds in the physical screening set. This is important because a focused library enables a more thorough evaluation of the anticancer potential of these molecules, using advanced physiologically relevant cancer models, incorporating genetically distinct patient-derived cell panel and complex pathological conditions (i.e.-host tumour microenvironment and hypoxia). With such a focused library, comprehensive screening paradigms including drug combinations, sequencing and scheduling can be incorporated into a primary phenotypic screening pipeline achievable by many academic screening facilities [53]. In the present work, we used fast, heuristic procedures to select the compounds with a predefined potency against the selected targets that presented with sufficient structural dissimilarity, as implemented in several filtering procedures. Even though our results show that the current heuristic approach leads to the desired results from the compound screening point of view, other approximate solutions to the MOP problem could further decrease the size of the screening set, while securing sufficient potency of the compounds against the targets.

Designing an optimal compound library of small molecules is challenged by the compound promiscuity, that is, many compounds modulate their effects through multiple protein targets with various degrees of potency. For instance, kinase inhibitors are notorious for their target promiscuity and their polypharmacological effects across various target classes beyond kinase families only [54]. The broad target selectivity of many compounds remain still uncharted [55], and therefore the phenotype driving targets of many compounds are currently unknown. In addition to the potency, chemical similarity of the compounds is another important feature of library design, especially if the aim is to have a collection of not too similar small-molecules that selectively target the protein space of interest. In the ideal case, each compound would selectively target only a single protein with high potency, leading to a diagonal design matrix between the compounds and proteins. However, the currently available small-molecule inhibitors remain quite far from this ideal scenario, hence requiring systematic design principles for library construction. Systematic study of cross-reactivity of the compounds will be needed to further investigate the target selectivity of the molecules in the current screening set across various target classes, beyond the rather limited data for target activity currently available for these compounds in public databases (**Suppl. Fig. 1D**). We are currently in the process of building and curating the C^3^L collection. Future applications in cell-based screening will provide important importation of the differential activity of the compounds in various cell-context, which can be used to further modify and tailor the library for more specific application cases.

Other future directions for the library construction include the use of phenotypic assays for biological profiling of compounds as cell-based features in the library design and analysis, including clustering of compounds into communities of phenotypic activity to explore biological mechanism of action (MoA) [56]. Further biological and clinical characterization, for instance, as implemented in the ChemicalChecker [57], could aid several drug discovery tasks, including target identification, MoA classification and library characterization. Integrated analysis of chemical, molecular target, cell-based profiling and clinical information could also be used to provide a relevance ranking of the targets and compounds in a library, once the disease indication is defined, hence adding one further dimension to the MOP problem. Other filtering steps to aid the eventual screening applications include, for instance, removal of interference compounds and finding on-target compounds with diverse scaffold profiles. The eventual applications in phenotypic screening, either in established cell lines or patient-derived cell models, will define the usefulness of any compound collections and libraries. We expect that the here-in presented comprehensive libraries with manually-curated target information will provide new opportunities for many exciting anticancer applications in GBM and other cancers, including target identification, drug repositioning and drug combination prediction and testing [58].

## Acknowledgements

The authors thank Dr. Drewry (UNC Eshelman School of Pharmacy) for his critical review and useful comments on the manuscript draft. The authors acknowledge funding support from Cancer Research UK and the Brain Tumour Charity (grant REF: C42454/A28596), and Helse Sør-Øst (grant No. 2020026).

## Supplementary Figures

**Suppl Figure 1.**
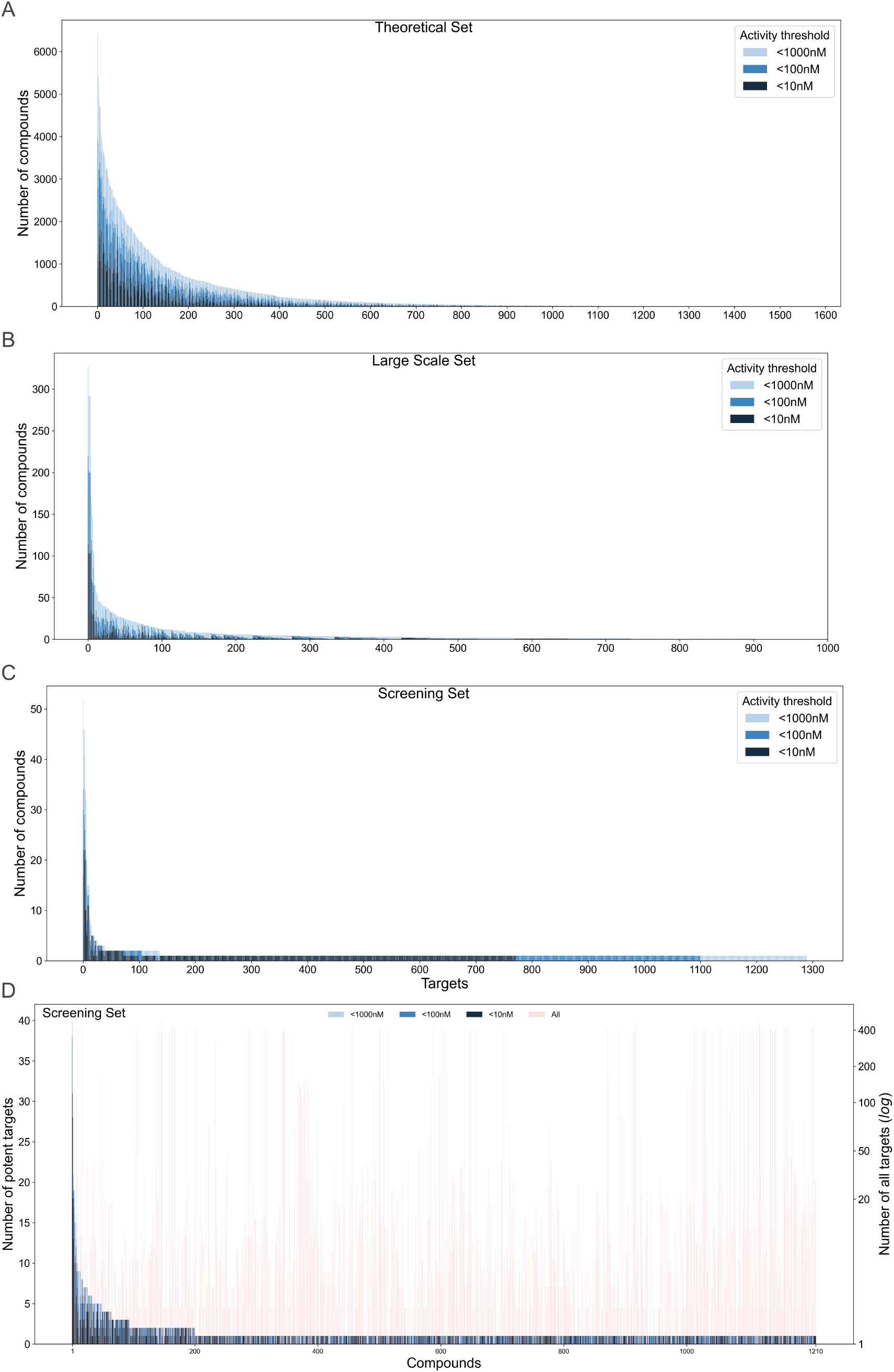
Number of compounds against targets using three activity thresholds. (A-C) Compounds in the three probe collections with different library sizes. Note the differences in the y-axis ranges. (D) Number of potent targets for the compounds in the Screening set using three activity thresholds. The right-hand y-axis depicts the total number of multi-dose bioactivity data for each compound-target pair available in Drug Target Commons, ChEMBL and DrugBank. Note the right-hand y-axis is log10-scaled while the numbers present the non-logged counts.

**Suppl Figure 2.**
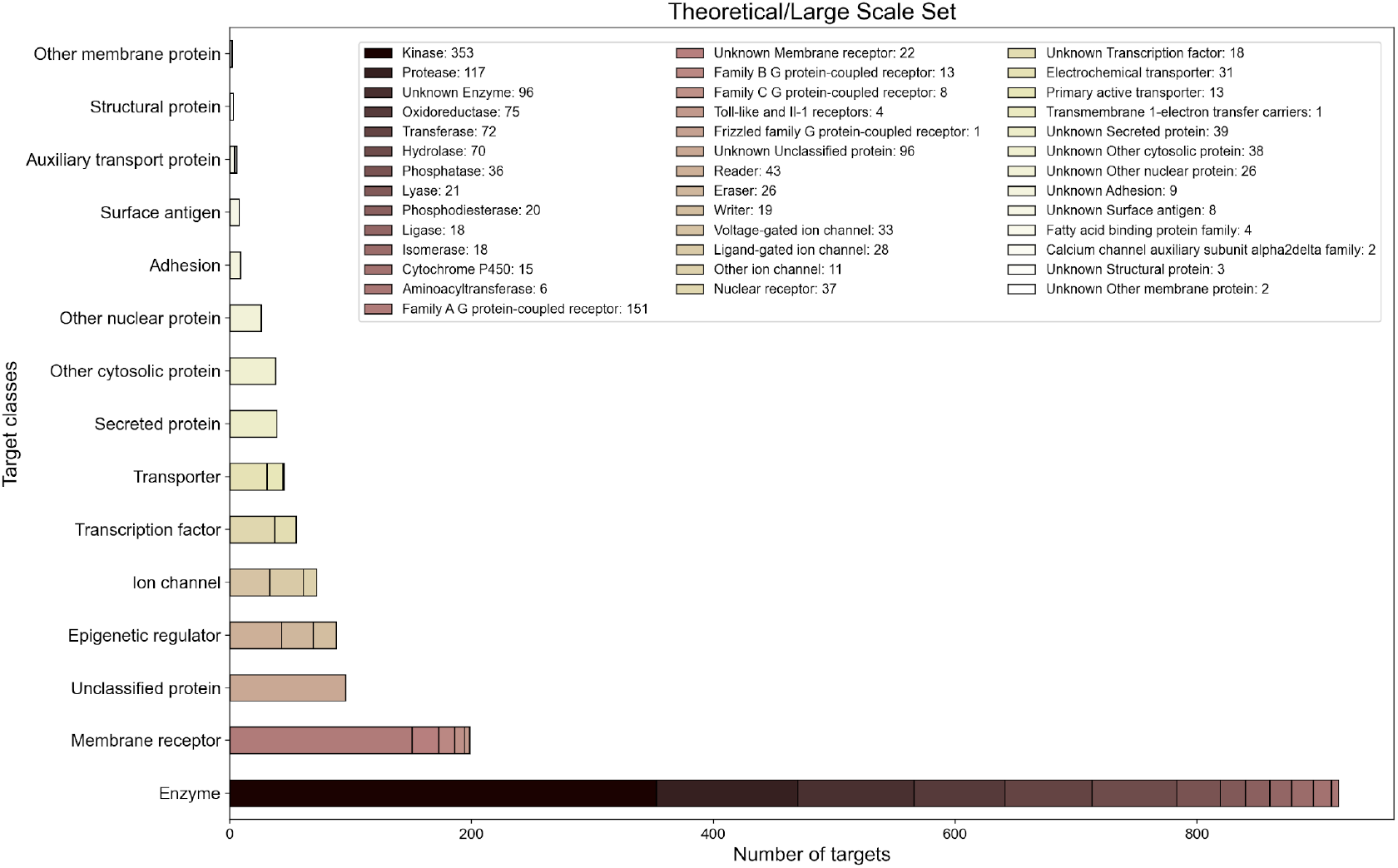
Target classes of the compounds in the Theoretical and Large-scale sets. The target classification of proteins was extracted from ChEMBL [19].

**Suppl Figure 3.**
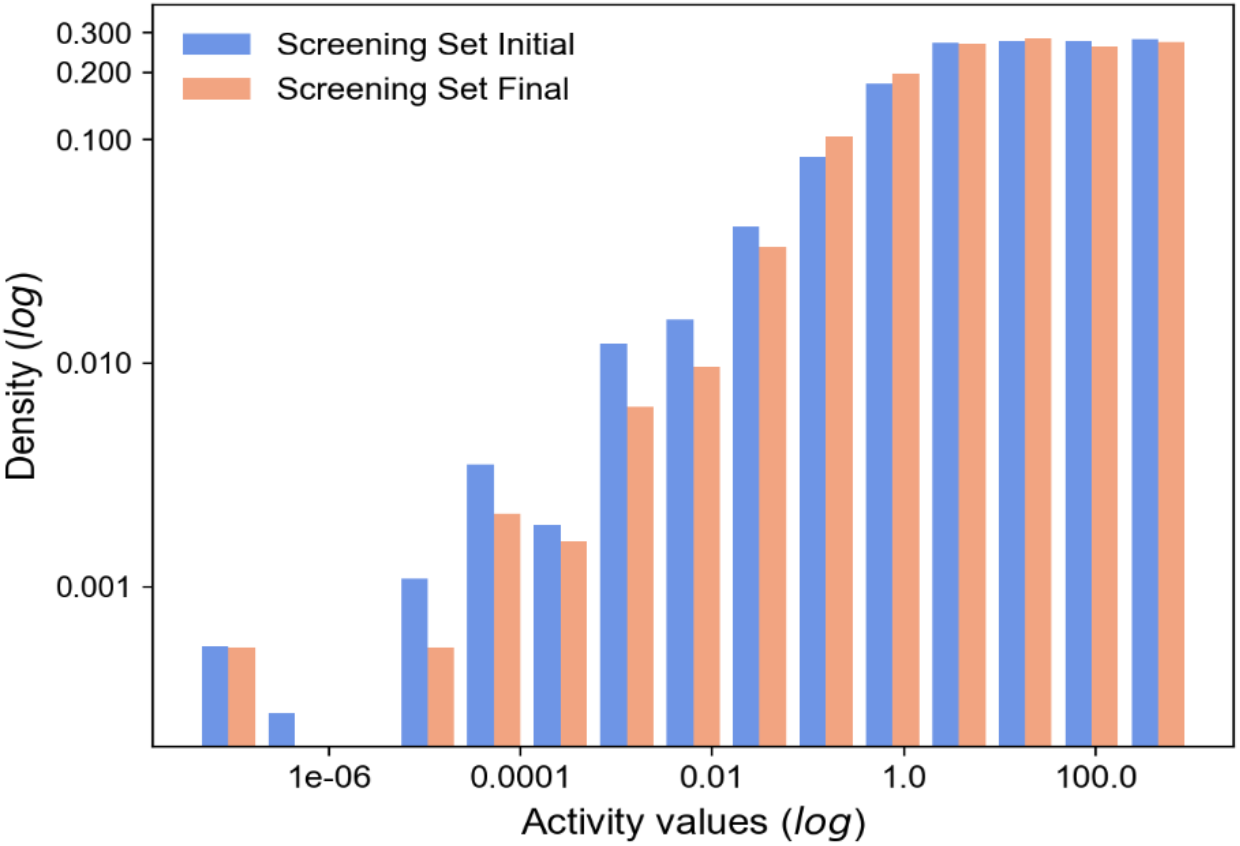
Comparison of the distributions of compound-target activities before and after replacing the unavailable compounds in the screening set. The x-axis and y-axis are log10-scaled, while the numbers present the non-logged values. The activity distributions were generally similar (p>0.05; Kolmogorov-Smirnov test).

## References

1. Doebele RC, Drilon A, Paz-Ares L, Siena S, Shaw AT, Farago AF, Blakely CM, Seto T, Cho BC, Tosi D, Besse B, Chawla SP, Bazhenova L, Krauss JC, Chae YK, Barve M, Garrido-Laguna I, Liu SV, Conkling P, John T, Fakih M, Sigal D, Loong HH, Buchschacher GL Jr, Garrido P, Nieva J, Steuer C, Overbeck TR, Bowles DW, Fox E, Riehl T, Chow-Maneval E, Simmons B, Cui N, Johnson A, Eng S, Wilson TR, Demetri GD; trial investigators. Entrectinib in patients with advanced or metastatic NTRK fusion-positive solid tumours: integrated analysis of three phase 1-2 trials. Lancet Oncol. 2020 Feb;21(2):271–282. doi: 10.1016/S1470-2045(19)30691-6. Epub 2019 Dec 11. Erratum in: Lancet Oncol. 2020 Feb;21(2):e70. Erratum in: Lancet Oncol. 2020 Jul;21(7):e341. Erratum in: Lancet Oncol. 2020 Aug;21(8):e372. PMID: 31838007; PMCID: PMC7461630.

2. Schultheis B, Strumberg D, Kuhlmann J, Wolf M, Link K, Seufferlein T, Kaufmann J, Feist M, Gebhardt F, Khan M, Stintzing S, Pelzer U. Safety, Efficacy and Pharcacokinetics of Targeted Therapy with The Liposomal RNA Interference Therapeutic Atu027 Combined with GemciTBine in Patients with Pancreatic Adenocarcinoma. A Randomized Phase Ib/IIa Study. Cancers (Basel). 2020 Oct 26;12(11):E3130. doi: 10.3390/cancers12113130. PMID: 33114652.

3. Zahavi D, Weiner L. Monoclonal Antibodies in Cancer Therapy. Antibodies (Basel). 2020 Jul 20;9(3):34. doi: 10.3390/antib9030034. PMID: 32698317; PMCID: PMC7551545.

4. van Zandwijk N, Pavlakis N, Kao SC, Linton A, Boyer MJ, Clarke S, Huynh Y, Chrzanowska A, Fulham MJ, Bailey DL, Cooper WA, Kritharides L, Ridley L, Pattison ST, MacDiarmid J, Brahmbhatt H, Reid G. Safety and activity of microRNA-loaded minicells in patients with recurrent malignant pleural mesothelioma: a first-in-man, phase 1, open-label, dose-escalation study. Lancet Oncol. 2017 Oct;18(10):1386–1396. doi: 10.1016/S1470-2045(17)30621-6. Epub 2017 Sep 1. PMID: 28870611.

5. Macedo N, Miller DM, Haq R, Kaufman HL. Clinical landscape of oncolytic virus research in 2020. J Immunother Cancer. 2020 Oct;8(2):e001486. doi: 10.1136/jitc-2020-001486. PMID: 33046622; PMCID: PMC7552841.

6. Robert A, Benoit-Vical F, Liu Y, Meunier B. Small Molecules: The Past or the Future in Drug Innovation? Metal Ions in Life Sciences. 2019 Jan;19. DOI: 10.1515/9783110527872-008.

7. van der Zanden SY, Luimstra JJ, Neefjes J, Borst J, Ovaa H. Opportunities for Small Molecules in Cancer Immunotherapy. Trends Immunol. 2020 Jun;41(6):493–511. doi: 10.1016/j.it.2020.04.004. Epub 2020 May 4. PMID: 32381382.

8. Pharmaceuticals B. Small Molecules to Treat Cancer - Research into Novel Therapies 2020 [updated Janueary 29, 2020. Available from: https://www.research.bayer.com/en/cancer-causes-cancer-research-new-drug-products-chemotherapy.aspx

9. Stupp R, Mason WP, van den Bent MJ, Weller M, Fisher B, Taphoorn MJ, Belanger K, Brandes AA, Marosi C, Bogdahn U, Curschmann J, Janzer RC, Ludwin SK, Gorlia T, Allgeier A, Lacombe D, Cairncross JG, Eisenhauer E, Mirimanoff RO; European Organisation for Research and Treatment of Cancer Brain Tumor and Radiotherapy Groups; National Cancer Institute of Canada Clinical Trials Group. Radiotherapy plus concomitant and adjuvant temozolomide for glioblastoma. N Engl J Med. 2005 Mar 10;352(10):987–96. doi: 10.1056/NEJMoa043330. PMID: 15758009.

10. Glioblastoma Stem Cells: Driving Resilience through Chaos, Prager, Briana C. et al. Trends in Cancer, Volume 6, Issue 3, 223–235

11. Horvath, P., Aulner, N., Bickle, M. et al. Screening out irrelevant cell-based models of disease. Nat Rev Drug Discov 15, 751–769 (2016). https://doi.org/10.1038/nrd.2016.175

12. US National Cancer Institute DDoCTD. Approved Oncology Drugs Set Information: A set of FDA-approved anticancer drugs to enable cancer research: National Institute of Health; 2020 [updated Sept 11, 2020. Available from: https://dtp.cancer.gov/organization/dscb/obtaining/available_plates.htm.

13. Chemicals S. Anti-cancer Compound Library 2020 [A unique collection of 3550 anti-cancer compounds for multiple cancers: Breast Cancer, Leukemia, Lung Cancer, Lymphoma, etc]. Available from: https://www.selleckchem.com/screening/anti-cancer-compound-library.html.

14. Patel, A., Seraia, E., Ebner, D., & Ryan, A. J. (2020). Adefovir dipivoxil induces DNA replication stress and augments ATR inhibitor-related cytotoxicity. International Journal of Cancer, 147(5), 1474–1484. https://doi.org/10.1002/ijc.32966

15. Hughes RE, Elliott RJR, Munro AF, et al. High-Content Phenotypic Profiling in Esophageal Adenocarcinoma Identifies Selectively Active Pharmacological Classes of Drugs for Repurposing and Chemical Starting Points for Novel Drug Discovery. SLAS Discovery: Advancing Life Sciences R & D. 2020 Aug;25(7):770–782. DOI: 10.1177/2472555220917115.

16. Hanahan D, Weinberg RA. Hallmarks of cancer: the next generation. Cell. 2011 Mar 4;144(5):646–74. doi: 10.1016/j.cell.2011.02.013. PMID: 21376230.

17. Wishart DS, Feunang YD, Guo AC, Lo EJ, Marcu A, Grant JR, Sajed T, Johnson D, Li C, Sayeeda Z, Assempour N, Iynkkaran I, Liu Y, Maciejewski A, Gale N, Wilson A, Chin L, Cummings R, Le D, Pon A, Knox C, Wilson M. DrugBank 5.0: a major update to the DrugBank daTBase for 2018. Nucleic Acids Res. 2017 Nov 8. doi: 10.1093/nar/gkx1037.

18. Smirnov, Petr, et al. “PharmacoDB: an integrative daTBase for mining in vitro anticancer drug screening studies.” Nucleic Acids Research (2017)

19. The ChEMBL daTBase in 2017, Gaulton A, Hersey A, Nowotka M, Bento AP, Chambers J, Mendez D, Mutowo P, Atkinson F, Bellis LJ, Cibrián-Uhalte E, Davies M, Dedman N, Karlsson A, Magariños MP, Overington JP, Papadatos G, Smit I, Leach AR. — Nucleic Acids Res. 2017; 45(D1):D945–D954. doi: 10.1093/nar/gkw1074

20. Drug Target Commons: A Community Effort to Build a Consensus Knowledge Base for Drug-Target Interactions Jing Tang, Zia-ur-Rehman Tanoli, Balaguru Ravikumar, Zaid Alam, Anni Rebane, Markus Vähä-Koskela, Gopal Peddinti, Arjan J. van Adrichem, Janica Wakkinen, Alok Jaiswal, Ella Karjalainen, Prson Gautam, Liye He, Elina Parri, Suleiman Khan, Abhishekh Gupta, Mehreen Ali, Laxman Yetukuri, Anna-Lena Gustavsson, Brinton Seashore-Ludlow, Anne Hersey, Andrew R. Leach, John P. Overington, Gretchen Repasky, Krister Wennerberg, Tero Aittokallio; Cell Chemical Biology, 2018,volume:25,pages:224–229

21. Forbes, S. A., Bhamra, G., Bamford, S., Dawson, E., Kok, C., Clements, J., Menzies, A., Teague, J. W., Futreal, P. A., & Stratton, M. R. (2008). The Catalogue of Somatic Mutations in Cancer (COSMIC). Current protocols in human genetics, Chapter 10, Unit– 10.11. https://doi.org/10.1002/0471142905.hg1011s57

22. Módos, D., Bulusu, K.C., Fazekas, D. et al. Neighbours of cancer-related proteins have key influence on pathogenesis and could increase the drug target space for anticancer therapies. npj Syst Biol Appl 3, 2 (2017). https://doi.org/10.1038/s41540-017-0003-6

23. Robert Roskoski, Properties of FDA-approved small molecule protein kinase inhibitors: A 2020 update, Pharmacological Research, Volume 152, 2020, doi: 10.1016/j.phrs.2019.104609

24. Stathias, V., Jermakowicz, A.M., Maloof, M.E. et al. Drug and disease signature integration identifies synergistic combinations in glioblastoma. Nat Commun 9, 5315 (2018). https://doi.org/10.1038/s41467-018-07659-z

25. Eid S., Turk S., Volkamer A., Rippmann F., and Fulle S. (2017) KinMap: a web-based tool for interactive navigation through human kinome data. BMC Bioinformatics, 18:16. DOI: 10.1186/s12859-016-1433-7

26. Denise Carvalho-Silva, Andrea Pierleoni, Miguel Pignatelli, ChuangKee Ong, Luca Fumis, Nikiforos Karamanis, Miguel Carmona, Adam Faulconbridge, Andrew Hercules, Elaine McAuley, Alfredo Miranda, Gareth Peat, Michaela Spitzer, Jeffrey Barrett, David G Hulcoop, Eliseo Papa, Gautier Koscielny, Ian Dunham, Open Targets Platform: new developments and updates two years on, Nucleic Acids Research, Volume 47, Issue D1, 08 January 2019, Pages D1056–D1065, https://doi.org/10.1093/nar/gky1133

27. Kim, S., Chen, J., Cheng, T., Gindulyte, A., He, J., He, S., Li, Q., Shoemaker, B. A., Thiessen, P. A., Yu, B., Zaslavsky, L., Zhang, J., & Bolton, E. E. (2019). PubChem 2019 update: improved access to chemical data. Nucleic acids research, 47(D1), D1102–D1109. https://doi.org/10.1093/nar/gky1033

28. R Development Core Team. R: A language and environment for statisticalcomputing.3-900051-07-0. R Foundation for Statistical Computing. Vienna,Austria, 2004.url:http://www.R-project.org

29. RDKit: Open-source cheminformatics; http://www.rdkit.org

30. Van Rossum G., & Drake, F. L. (2009). Python 3 Reference Manual. Scotts Valley, CA: CreateSpace

31. The Human Protein Atlas, Uhlén M, Fagerberg L, Hallström BM, Lindskog C, Oksvold P, Mardinoglu A, Sivertsson Å, Kampf C, Sjöstedt E, Asplund A, Olsson I, Edlund K, Lundberg E, Navani S, Szigyarto CA, Odeberg J, Djureinovic D, Takanen JO, Hober S, Alm T, Edqvist PH, Berling H, Tegel H, Mulder J, Rockberg J, Nilsson P, Schwenk JM, Hamsten M, von Feilitzen K, Forsberg M, Persson L, Johansson F, Zwahlen M, von Heijne G, Nielsen J, Pontén F. Tissue-based map of the human proteome. Science 2015 347(6220):1260419. PubMed: 25613900 DOI: 10.1126/science.1260419

32. Wanjuan Yang, Jorge Soares, Patricia Greninger, Elena J. Edelman, Howard Lightfoot, Simon Forbes, Nidhi Bindal, Dave Beare, James A. Smith, I. Richard Thompson, Sridhar Ramaswamy P. Andrew Futreal, Daniel A. Haber, Michael R. Stratton Cyril Benes, Ultan McDermott, Mathew J. Garnett, Genomics of Drug Sensitivity in Cancer (GDSC): a resource for therapeutic biomarker discovery in cancer cells, Nucleic Acids Research, Volume 41, Issue D1, 1 January 2013, Pages D955–D961, https://doi.org/10.1093/nar/gks1111

33. Correlating chemical sensitivity and basal gene expression reveals mechanism of action” Rees et al., Nat Chem Biol, 12, 109-116 (2016), and “Harnessing Connectivity in a Large-Scale Small-Molecule Sensitivity Dataset” Seashore-Ludlow et al., Cancer Discovery, 5, 1210-1223 (2015), and “An Interactive Resource to Identify Cancer Genetic and Lineage Dependencies Targeted by Small Molecules” Basu, Bodycombe, Cheah, et al., Cell, 154, 1151–1161 (2013)

34. Barretina, J., Caponigro, G., Stransky, N. et al. The Cancer Cell Line Encyclopedia enables predictive modelling of anticancer drug sensitivity. Nature 483, 603–607 (2012). https://doi.org/10.1038/nature11003

35. Schönbach, C., Koh, J. L., Flower, D. R., Wong, L., & Brusic, V. (2002). FIMM, a daTBase of functional molecular immunology: update 2002. Nucleic acids research, 30(1), 226–229. https://doi.org/10.1093/nar/30.1.226

36. Daemen, A., Griffith, O.L., Heiser, L.M. et al. Modeling precision treatment of breast cancer. Genome Biol 14, R110 (2013). https://doi.org/10.1186/gb-2013-14-10-r110, and Laura M. Heiser, Anguraj Sadanandam, Joe W. Gray, et al. “Subtype and pathway specific responses to anticancer compounds in breast cancer” Proceedings of the National Academy of Sciences Feb 2012, 109 (8) 2724–2729; DOI: 10.1073/pnas.1018854108

37. Jaiswal A, Yadav B, Wennerberg K, Aittokallio T. Integrated Analysis of Drug Sensitivity and Selectivity to Predict Synergistic Drug Combinations and Target Coaddictions in Cancer. Methods Mol Biol. 2019;1888:205–217. doi: 10.1007/978-1-4939-8891-4_12. PMID: 30519949.

38. Fazekas, D. et al. SignaLink 2—a signaling pathway resource with multi-layered regulatory networks. BMC. Syst. Biol. 7, 7 (2013)

39. Croft, D. et al. The Reactome pathway knowledgebase. Nucleic. Acids. Res. 42, D472–D477 (2014)

40. Keshava Prasad T. S. et al. Human protein reference daTBase--2009 update. Nucleic. Acids. Res. 37, D767–D772 (2009)

41. Salwinski, L. et al. The daTBase of interacting proteins: 2004 update. Nucleic. Acids. Res. 32, D449–D451 (2004)

42. Orchard, S. et al. The MIntAct project--IntAct as a common curation platform for 11 molecular interaction daTBases. Nucleic. Acids. Res. 42, D358–D363 (2014)

43. Stark, C. et al. BioGRID: a general repository for interaction datasets. Nucleic. Acids. Res. 34, D535–D539 (2006)

44. Cui, Q. et al. A map of human cancer signaling. Mol. Syst. Biol. 3, 152 (2007)

45. Bozdogan, H. (1987). Model selection and Akaike’s Information Criterion (AIC): The general theory and its analytical extensions. Psychometrika, 52(3), 345–370. doi:10.1007/bf02294361

46. Irwin JJ, Shoichet BK. ZINC--a free daTBase of commercially available compounds for virtual screening. J Chem Inf Model. 2005 Jan-Feb;45(1):177–82. doi: 10.1021/ci049714+. PMID: 15667143; PMCID: PMC1360656.

47. Bailey MH, Tokheim C, Porta-Pardo E, Sengupta S, Bertrand D, Weerasinghe A, Colaprico A, Wendl MC, Kim J, Reardon B, Ng PK, Jeong KJ, Cao S, Wang Z, Gao J, Gao Q, Wang F, Liu EM, Mularoni L, Rubio-Perez C, Nagarajan N, Cortés-Ciriano I, Zhou DC, Liang WW, Hess JM, Yellapantula VD, Tamborero D, Gonzalez-Perez A, Suphavilai C, Ko JY, Khurana E, Park PJ, Van Allen EM, Liang H; MC3 Working Group; Cancer Genome Atlas Research Network, Lawrence MS, Godzik A, Lopez-Bigas N, Stuart J, Wheeler D, Getz G, Chen K, Lazar AJ, Mills GB, Karchin R, Ding L. Comprehensive Characterization of Cancer Driver Genes and Mutations. Cell. 2018 Apr 5;173(2):371–385.e18. doi: 10.1016/j.cell.2018.02.060. Erratum in: Cell. 2018 Aug 9;174(4):1034-1035. PMID: 29625053; PMCID: PMC6029450

48. ICGC/TCGA Pan-Cancer Analysis of Whole Genomes Consortium. Pan-cancer analysis of whole genomes. Nature. 2020 Feb;578(7793):82–93. doi: 10.1038/s41586-020-1969-6. Epub 2020 Feb 5. PMID: 32025007; PMCID: PMC7025898.

49. Ravikumar B, Timonen S, Alam Z, Parri E, Wennerberg K, Aittokallio T. Chemogenomic Analysis of the Druggable Kinome and Its Application to Repositioning and Lead Identification Studies. Cell Chem Biol. 2019 Nov 21;26(11):1608–1622.e6. doi: 10.1016/j.chembiol.2019.08.007. Epub 2019 Sep 11. PMID: 31521622.

50. Gautam P, Jaiswal A, Aittokallio T, Al-Ali H, Wennerberg K. Phenotypic Screening Combined with Machine Learning for Efficient Identification of Breast Cancer-Selective Therapeutic Targets. Cell Chem Biol. 2019 Jul 18;26(7):970–979.e4. doi: 10.1016/j.chembiol.2019.03.011. Epub 2019 May 2. PMID: 31056464; PMCID: PMC6642004.

51. Elkins JM, Fedele V, Szklarz M, Abdul Azeez KR, Salah E, Mikolajczyk J, Romanov S, Sepetov N, Huang XP, Roth BL, Al Haj Zen A, Fourches D, Muratov E, Tropsha A, Morris J, Teicher BA, Kunkel M, Polley E, Lackey KE, Atkinson FL, Overington JP, Bamborough P, Müller S, Price DJ, Willson TM, Drewry DH, Knapp S, Zuercher WJ. Comprehensive characterization of the Published Kinase Inhibitor Set. Nat Biotechnol. 2016 Jan;34(1):95–103. doi: 10.1038/nbt.3374. Epub 2015 Oct 26. PMID: 26501955.

52. Corsello SM, Bittker JA, Liu Z, Gould J, McCarren P, Hirschman JE, Johnston SE, Vrcic A, Wong B, Khan M, Asiedu J, Narayan R, Mader CC, Subramanian A, Golub TR. The Drug Repurposing Hub: a next-generation drug library and information resource. Nat Med. 2017 Apr 7;23(4):405–408. doi: 10.1038/nm.4306. PMID: 28388612; PMCID: PMC5568558.

53. Shanks, E., Ketteler, R. & Ebner, D. Academic drug discovery within the United Kingdom: a reassessment. Nat Rev Drug Discov 14, 510 (2015). https://doi.org/10.1038/nrd4661

54. Moret N, Clark NA, Hafner M, Wang Y, Lounkine E, Medvedovic M, Wang J, Gray N, Jenkins J, Sorger PK. Cheminformatics Tools for Analyzing and Designing Optimized Small-Molecule Collections and Libraries. Cell Chem Biol. 2019 May 16;26(5):765–777.e3. doi: 10.1016/j.chembiol.2019.02.018. Epub 2019 Apr 4. PMID: 30956147; PMCID: PMC6526536.

55. Santos R, Ursu O, Gaulton A, Bento AP, Donadi RS, Bologa CG, Karlsson A, Al-Lazikani B, Hersey A, Oprea TI, Overington JP. A comprehensive map of molecular drug targets. Nat Rev Drug Discov. 2017 Jan;16(1):19–34. doi: 10.1038/nrd.2016.230. Epub 2016 Dec 2. PMID: 27910877; PMCID: PMC6314433.

56. Dančík V, Carrel H, Bodycombe NE, Seiler KP, Fomina-Yadlin D, Kubicek ST, Hartwell K, Shamji AF, Wagner BK, Clemons PA. Connecting Small Molecules with Similar Assay Performance Profiles Leads to New Biological Hypotheses. J Biomol Screen. 2014 Jun;19(5):771–81. doi: 10.1177/1087057113520226. Epub 2014 Jan 24. PMID: 24464433; PMCID: PMC5554958.

57. Duran-Frigola M, Pauls E, Guitart-Pla O, Bertoni M, Alcalde V, Amat D, Juan-Blanco T, Aloy P. Publisher Correction: Extending the small-molecule similarity principle to all levels of biology with the Chemical Checker. Nat Biotechnol. 2020 Sep;38(9):1098. doi: 10.1038/s41587-020-0564-6. Erratum for: Nat Biotechnol. 2020 Sep;38(9):1087–1096. PMID: 32440008.

58. Tarasov VV, Svistunov AA, Chubarev VN, Zatsepilova TA, Preferanskaya NG, Stepanova OI, Sokolov AV, Dostdar SA, Minyaeva NN, Neganova ME, Klochkov SG, Mikhaleva LM, Somasundaram SG, Kirkland CE, Aliev G. Feasibility of Targeting Glioblastoma Stem Cells: From Concept to Clinical Trials. Curr Top Med Chem. 2019;19(32):2974–2984. doi: 10.2174/1568026619666191112140939. PMID: 31721715.

